# Whole-genome sequencing and phenotyping reveal specific adaptations of *Lachancea thermotolerans* to the winemaking environment

**DOI:** 10.1101/2024.05.31.596785

**Authors:** Javier Vicente, Anne Friedrich, Joseph Schacherer, Kelle Freel, Domingo Marquina, Antonio Santos

## Abstract

Adaptation to the environment plays an essential role in yeast evolution as a consequence of selective pressures. *Lachancea thermotolerans*, a yeast related to the fermentative process and one of the current trends in wine technology research, has undergone an anthropization process that has led to a strong differentiation both from a genomic and phenomic perspective. Using whole-genome sequencing, we have investigated the genomic diversity of 145 *L. thermotolerans* strains, identifying six well-defined groups primarily delineated by their ecological origin and exhibiting high levels of genetic diversity. Anthropized strains showed lower genetic diversity due to the purifying selection imposed by the winemaking environment. Strong evidence of anthropization and adaptation to the wine environment through modification of gene content was also found. Differences in genes involved in the assimilation of alternative carbon and nitrogen sources, such as the *MAL1* and *DAL5* genes, which confer greater fitness in the winemaking environment, were observed. Additionally, we found that phenotypic traits considered domestication hallmarks are present in anthropized strains. Among these, increased fitness in the presence of ethanol and sulphites, assimilation of non-fermentable carbon sources such as glycerol, and lower levels of residual fructose under fermentative conditions highlight. We hypothesize that lactic acid production by wine-related strains is an anthropization signature consequence of the adaption of early Crabtree-positive yeasts to the fermentative environment. Overall, the results of this work provide valuable insight into the anthropization process in *L. thermotolerans* and demonstrate how fermentation environments give rise to similar adaptations in different yeast species.

## INTRODUCTION

Adaptation to anthropic niches has been linked to different genomic and phenotypic changes arising from several traits of industrial relevance. Usually, most definitions of the domestication process are anthropocentric and centred on human intentionally, without taking into consideration an unintentional selection. Microbial adaptation to different industrial processes is traditionally consequence of this accidental selection (Friedrich et al., 2023). The ubiquitous presence of microorganisms and their importance in different human-related processes has been one of the main drivers of microbial adaptation and specialization (Villarreal et al., 2022).

Detecting the selective pressures driving genomic variation is essential to better understand adaptation and predict how natural populations will evolve in response to changing environmental conditions. Environmental factors generate differential signatures on the genomes of species and populations, with biotic and abiotic influences often acting at different levels and affecting various genomic features. Studies developed using whole genomes can improve the identification of selective pressures driving niche-specific adaptation patterns in different organisms and environments (Dauphin et al., 2023). The different environments in which the species evolved led to the emergence of specific phenotypic traits associated with each subpopulation, ultimately resulting in a strong genetic differentiation and a robust population structure (Caudal et al., 2022; Peris et al., 2023; Peter et al., 2018; Teyssonniere et al., 2024).

Wine production, which is probably the most valuable fermented beverage due to its cultural, religious, and economic importance, is far to be a single species process (García-Ríos & Guillamón, 2019). During wine fermentation, several yeasts species and genera inhabiting the grape surface and winery dependencies, participate in a competition for resource and stresses survival. Microorganisms inhabiting grape must face several challenging conditions such as high osmolarity, low pH, depletion of different nutritional factors (e.g., nitrogen, lipids, or vitamins) and the presence of diverse antimicrobials (e.g., ethanol and sulphites). These factors act either individually or synergistically, leading to adaptative differentiation (Marsit & Dequin, 2015). *Saccharomyces cerevisiae,* not only strongly associated to this environment, but also with other anthropic niches such as beer, bread, or sake, is the most studied with regard to genomics and phenotypic diversity. Exploration of thousands of genomes over the past few years has revealed different independent and linage-specific domestication events within the species (Ohnuki et al., 2017; Peter et al., 2018; Saada et al., 2023; Schacherer et al., 2009). These events led to different evolutionary trajectories and the emergence of unique metabolic traits that have made *S. cerevisiae* the model species on yeasts population genomics and niche-adaptation studies (Belda et al., 2019; Camarasa et al., 2011; Gallone et al., 2016; Marr et al., 2023; Peris et al., 2023; Peter et al., 2018; Peter et al., 2022; Ruiz et al., 2021; Warringer et al., 2011).

Despite the interest in *S. cerevisiae* as a model species in evolutionary genomics, recent studies have also focused on other non-*Saccharomyces* species participating in wine fermentations. These include *Torulaspora delbrueckki* (Silva et al., 2023), *Brettanomyces bruxellensis* (Eberlein et al., 2021; Gounot et al., 2020), *Hanseniaspora uvarum* (Albertin et al., 2016), and *Lachancea thermotolerans (Freel et al., 2014)* (Hranilovic et al., 2017). Other yeasts genera associated with different industrial processes, such as *Kluyveromyces lactis* (Friedrich et al., 2023; Varela et al., 2019) or *Yarrowia lypolitica* (Bigey et al., 2023), as well as those of ecological significance as *Saccharomyces uvarum* (Almeida 2014), *Lachancea cidri* (Villarreal et al., 2024; Villarreal et al., 2022), and *Schizosaccharomyces pombe* (Jeffares et al., 2015) have gained attention. Notably, yeasts species of clinical relevance such as *Candida glabrata* (Wang et al., 2024), *Candida auris* (Chow et al., 2020) or *Cryptococcus neoformans* (Sephton-Clark et al., 2022), are also being studied. Broadening the areas of research could provide a better understanding of the evolutionary dynamics of yeasts (and fungi) and highlight the ecological importance of their intraspecific diversity. Access to whole genome sequencing data from large numbers of individuals undergoing adaptation processes provides valuable information on the genomic footprints of adaptation throughout the entire evolutionary tree (Friedrich et al., 2023; Peter & Schacherer, 2016).

The yeast species *L. thermotolerans*, a non-*Saccharomyces* yeast involved in wine fermentation, has recently attracted attention for its lactic acid production during alcoholic fermentation. This characteristic is considered a valuable approach in winemaking as it provides a biological approach to deal with the effects of climate change on grape musts, such as a higher concentration of fermentable sugars and a decrease in acidity values (Hranilovic et al., 2018; Porter et al., 2019; Vicente et al., 2023; Vilela, 2018). Additionally, several *L. thermotolerans* enzymes have been studied due to their potential in industrial and pharmaceutical applications (Aktar et al., 2023; Fan et al., 2020; Mousavi et al., 2015; Prasad et al., 2005).

*L. thermotolerans* is the type species of the genus *Lachancea,* initially included within the *Kluyveromyces* clade (Lachance & Kurtzman, 2011). This genus diverged after the appearance of anaerobic capability, estimated to have occurred around 125-150 million years ago, prior to the whole-genome duplication event approximately 100 million years ago. This clade represents the first lineage after the loss of respiratory chain complex I, which happened after the split of the *Saccharomyces*-*Lachancea* and *Kluyveromyces*-*Eremothecium* lineages around 125-150 million years ago. This event allowed the emergence of the long-term Crabtree effect and the ability to grow under anaerobic conditions (Dashko et al., 2014; Hagman et al., 2014; Motlhalamme et al., 2022). Therefore, focusing on this genus, as well as in individual species within it, could be crucial for understanding the origin of one of the most important traits related to carbon metabolism in yeasts and inferring the earliest molecular events initiating Crabtree effect in the most ancestral species (Hagman et al., 2013).

In terms of population genomics, species within the *Lachancea* clade show great diversity compared to those in the *Saccharomyces* clade (Villarreal et al., 2022). The different species conforming the genus have been found in different environments, ranging from natural habitats to anthropized ones, including tree barks and exudates, flowers, insects, soils, food, and beverages. Despite its ubiquitous presence, information regarding genetic adaptations to these environments and associated phenotypic traits remains limited. To date, although the studies on *L. thermotolerans* regarding its differentiation in terms of genetic and phenotypic diversity, only the population genomics and adaptive processes of *L. cidri* have been carefully addressed.

The intraspecific diversity of *L. thermotolerans* has been previously addressed by different approaches, including mitochondrial genomics and microsatellite studies (Banilas et al., 2016; Freel et al., 2014; Hranilovic et al., 2017). These studies highlighted the role of geographic isolation and local adaptation as drivers of the evolutionary process in this species. The first work on *L. thermotolerans* intraspecific diversity involved the analysis of mitochondrial DNA restriction patterns in a limited number of strains, revealing a low level of intraspecific divergence and suggesting the influence of different evolutionary forces (Belloch et al., 1997). Further studies comprising a higher number of strains and higher-resolution techniques corroborated this fact. They demonstrated that ecological niche (Freel et al., 2014) and geographical location (Banilas et al., 2016) may act as driving forces differentiating subpopulations within the species, with the oenological environment playing an essential role in this divergence (Hranilovic et al., 2017). Notably, significant phenotypic differences resulting from the anthropization process, in particular adaptation to the winemaking environment, were observed, with these isolates exhibiting traits relevant for wine production (Hranilovic et al., 2018). Furthermore, subsequent studies using whole genome sequencing have revealed significant genetic differences between high and low lactic acid producing strains (Gatto et al., 2020).

However, the population structure and gene content, as well as the genetic variation underlying the phenotypic diversity of this species, have not yet been addressed. To have a better insight into these two facets, we characterized the genomic and phenotypic profiles by whole genome sequencing and deep phenotyping of 145 *L. thermotolerans* strains of diverse geographical and ecological origins. *L. thermotolerans* CBS 6340 has been previously sequenced and annotated (Souciet et al., 2009) and was therefore used as reference for population genomic sand gene content analysis. Exploring the overall pattern of polymorphism allowed us to stablish precise phylogenetic relationship among strains, revealing different subpopulations composed of closely related isolates. By identifying CNVs (Copy Number Variants) and studying the variation in gene content, we detected several genes, related to adaptation and specialization to the winemaking environment, as well as different phenotypes associated with anthropization and wine-related traits. The fermentation performance of the sequenced strains not only agreed with the phylogenetic results, but also revealed that important traits, notably in winemaking and the use of this species as a biological tool for acidity management, were influenced by the anthropization process.

## MATERIAL AND METHODS

### Yeast strains, DNA extraction and sequencing

The collection of 145 *L. thermotolerans* strains from diverse ecological and geographical origins included in this study is listed in Table S1. The collection is composed by strains from all continents, excluding Antarctica, isolated in diverse ecological niches, although almost half of the isolates were obtained from anthropized environments, mainly wine-related ones.

All samples underwent Illumina paired-end sequencing, which was carried out in two separate batches (as indicated in Table S1). For the first batch (1), a single colony was inoculated in 10 mL of YNB-glucose 2% (Difco) using 50 mL flasks and cultured at 25°C and 150 rpm for 20 hours. Then, yeast cells were centrifuged and washed using saline solution. Genomic DNA was extracted using the QIAGEN Genomic-tip 20/G isolation kit, and Illumina sequencing libraries were prepared using the TruSeq Nano DNA Kit. The libraries were then subjected to paired-end sequencing (2×150) on an Illumina NovaSeq 6000 sequencer. For the second batch (2), yeast cells were cultured overnight at 30°C in 20 mL of YPD medium until reached the early stationary phase, then cells were harvested by centrifugation and total genomic DNA was extracted using the Masterpure Yeast DNA purification kit. Genomic Illumina sequencing libraries were prepared, multiplexed and subjected to pair-end sequencing (2×100) using a HiSeq 2000 sequencer.

### Mapping, variant calling and quality filtering

For each sample, Illumina paired-end sequences were aligned to the reference genome of *L. thermotolerans* CBS 6340 (https://forgemia.inra.fr/gryc-data) (Souciet et al., 2009) using BWA mem (v0.7.17-r1188) with default parameters (Li & Durbin, 2009). Alignments were sorted using samtools (v1.6), processed with GATK (v4.3.0.0) AddOrReplaceReadGroups and indexed using samtools (Danecek et al., 2021; McKenna et al., 2010). SNPs (Single Nucleotide Polymorphisms) and indels (Insertion-Deletion mutations) detection as well as genotyping were conducted simultaneously across all samples using the GATK pipeline (HaplotypeCaller, GenomicsDBImport, and GenotypeGVCFs). The identified SNPs and indels were then segregated into separate files, and quality filters were applied using SelectVariants and VariantFiltration options from GATK with specific parameter thresholds (QD < 2.0, QUAL < 30.0, SOR > 3.0, FS > 60.0, MQ < 40.0, MQRankSum < -12.5, ReadPosRankSum < -8.0).

### Population genomics: tree building, population structure and genetics

The phylogeographic relationships between samples was determined from a filtered SNP file resulting from the variant calling analysis. The tree was constructed according to the BIONJ algorithm and bootsrap = 100 in R (v4.3.1) using the SNPRelate (v1.34.1), ape (v5.7-1), pvclust (v2.2-0) and ggtree packages (v3.8.0).

Population structure and individual admixture proportions were estimated from genotype likelihoods of biallelic SNPs using FastStructure (v1.0) and ADMIXTURE (v1.3.0) (Alexander et al., 2009; Pritchard et al., 2000). The number of ancestral populations (K) was tested from 2 to 8 to estimate individual ancestral proportions for each structure model using default parameter values for stop criteria and SNP filtering. Plots with individual ancestries were constructed with R using ggplot2 package (v3.4.2).

Pairwise nucleotide diversity (π) and Tajima’s D value were calculated using VCFtools (v0.1.16) (Danecek et al., 2011) both for the whole population and for the different genetic clusters. Fixation-indexes (F_ST_) across genomes were calculated using SNPRelate (v1.34.1) and vcfR (v1.14.0) R packages for the different genetic clusters.

### Pangenome and gene content variation

The study of the pangenome was performed through two approaches. Genes present in the reference strain were analysed using control-FREEC (v11.4) (Boeva et al., 2012) (window = 1; breakPointThreshold = 0.05; minExpectedGC = 0.35; maxExpectedGC = 0.55; ploidy = 2), that allowed at the same time the determination of CNVs (Copy Number Variants) in each sample. The set of genes not present in the reference strain, but present in any of the strains of the dataset, was determined using a previously described pipeline (Gounot et al., 2020). In brief, assemblies were constructed using SPADES (v3.13.0) (Prjibelski et al., 2020) with 127 and 63 as kmer sizes depending on the sequencing technique of the strains (2×150 or 1×100). The assemblies were compared against the reference using BlastN, allowing the determination of non-reference fragments. Gene prediction of these fragments was based on SNAP and Augustus with the training sets generated using the *L. thermotolerans* CBS 6340 reference proteome. Genes showing low complexity regions were discarded, as well as those whose product had less than 50 amino acids. All genes were then compared to each other using BlastP, employing a graph-based method to identify the non-redundant set of non-reference fragments.

Once the entire dataset of genes was retrieved, they were classified into core and supplemental genes, for those present in all the studied dataset or for those absent in any of the studied strains respectively. This classification was further subdivided into soft-core genome (genes present in at least the 95 % of the strains), shell genome (genes present in between 95 to 5 %) and cloud genome (genes present in less than 5 % of the strains), as previously described (Contreras-Moreira & Vinuesa, 2013; Silva et al., 2023). All cloud genes, prone to be biased by sequencing and prediction strategies, were then compared against the NCBI non-redundant *Saccharomycetales* database using BlastP, and those showing no significant hits (e-value < 1e-4) were discarded. All non-reference genes were annotated through similarity searches (e-value < 1e-4) using the eggnog (v2.1.12) database (Huerta-Cepas et al., 2019).

The expected pangenome saturation curves were calculated using the specaccum function of the vegan package (v2.6-4). The openness of the pangenome was modelled using 1,000 permutations according to Heap’s law (n = kN^α^; where n is the pan-genome size, N is the number of genomes used, and k and α are the fitting parameters) (Tettelin et al, 2014), using R package micropan (v2.1).

### Phylogenetic reconstruction of the species

To construct the phylogenetic tree of the species, we employed 20 of our sequenced strains together with a strain representative of each species in the *Lachancea* genera and used *Kluyveromyces lactis* as an outgroup. First, we determined the common set of orthologous genes using the reference genomes of all the species of interest downloaded from (https://forgemia.inra.fr/gryc-data). Orthologs were determined using Proteinortho (v6.0.18) and default parameters (-p=blastn option) (Lechner et al., 2011). 153 one-to-one orthologs with a minimum algebraic connectivity of 0.1 were selected for further analysis. BLAST was employed to extract the orthologous from our sequenced *L. thermotolerans* strains using the sequence from the reference strain as query. All orthologs were concatenated by sample aligned with MAFFT (v7.520) using default parameter values (Katoh & Standley, 2013). Maximum likelihood tree was constructed using 100 iterations and visualized using the ggtree R package (v3.8.0).

### *L. thermotolerans* high throughput phenotyping

The phenotypic characterization of the yeasts was conducted by measuring different growth parameters in 25 different conditions, encompassing 11 fermentation conditions in Synthetic Grape Must (SGM) and the use of 14 different carbon and nitrogen sources at different temperatures (described in Table S2).

SGM control medium was prepared as previously described (Henschke & Jiranek, 1993) adjusting final Yeast Assimilable Nitrogen (YAN) to 200 mg N/L (60 mg N/L of ammonia-nitrogen ((NH_4_)_2_HPO_4_) and 140 mg N/L of amino acids). The growth capacity in different carbon sources was assayed in Synthetic Medium (SM) prepared with Yeast Nitrogen Base 0.67% (BD Difco™, USA), and 2.0 % of the corresponding carbon source. Likewise, nitrogen source conditions were assessed in SM prepared with Yeast Carbon Base 0.67% (BD Difco™, USA), and 100 mg N/L of the corresponding nitrogen source. For growth assays at different temperatures SM-glucose was employed. All media were sterilised by 0.22 μm filtration.

Yeast strains were pre-cultivated in 96-well plates for 10 hours in 250 μL of the corresponding control media for each assay. Then, precultures were diluted 1 to 100 in saline solution (NaCl 0.9%) and 10 μL were inoculated by triplicate in 240 μL of the corresponding media. All cultures transfers and inoculations were performed employing a robotic system (Assist Plus – VIALAB system, Integra Bioscience, Switzerland). Initial optical density at 600 nm (OD_600_) of all cultures were between 0.03-0.05. Assays were conducted at 25 °C (except for those conditions where temperature was tested as a variable), with orbital shaking at 100 rpm. Culture growth was monitored by measuring OD_600_ (with a previous orbital agitation of plates at 300 rpm for 5 seconds) at different time points during 70 hours of culture, using a microplate reader (Varioskan Flash Multimode Reader, Thermo Scientific, USA). Then, measured data were adjusted to a Baranyi growth model using the R package GrowthRates (v0.8.4) and the different growth parameters (lag phase, growth rate and proliferative efficiency) were extracted.

### Fermentative performance of the studied strains

The fermentative performance of the studied strains was assayed in SGM prepared as described previously. Yeast were precultured in 2 ml of SGM in 24-well plates for 24 hours at 25°C and 100 rpm. Subsequently, 0.5 mL of the resuspended precultures were inoculated in triplicate into 14 mL of SGM. Fermentations were carried out at 25°C for 7 days (168 hours). Afterward, cultures were centrifugated and stored at -70°C for further analysis. Different oenological parameters (ethanol, residual sugars, glycerol, total acidy, pH, tartaric, citric, malic, lactic, acetic, and succinic acids) were determined by Fourier-transform infrared spectroscopy (Bachus 3 MultiSpec, *Tecnología Difusión Ibérica*, S.L, Spain).

### Correlation between phylogeographic relationships and phenotype

To determine the Pearson’s correlation between the phylogeographic distribution of the studied strains with the phenotyping data, the phylogenetic distance between strains was calculated as the sum of branch lengths between each pair. Meanwhile, the phenotypic distance was calculated as the Euclidean distance between strains based on the phenotypic data, including lag phase and maximum growth rate obtained from high throughput phenotyping, as well as oenological parameters from the fermentative performance of the strains. For each trait, the Pagel’s λ value was determined as previously described (Ruiz et al., 2023). All calculations were made using ape (v5.7-1) and phytools (v2.0-3) R packages.

## RESULTS

### The evolutionary history of *L. thermotolerans* is defined by the wine environment

The genomes of 145 different strains of *L. thermotolerans* isolated in different regions (Africa, South America, North America, Asia, Australia, and Europe) and substrates (winemaking, plant, animals) were sequenced, with a mean coverage of 112-fold. For each sample, reads were mapped to the reference sequence (*L. thermotolerans* CBS 6340), allowing the detection of a total of 1,008,631 reference-based polymorphic positions of which 947,516 were biallelic SNPs and 39,294 small indels. All sequenced genomes are euploid, with no evidence of aneuploidy.

The resulting set of 947,516 biallelic SNPs was used to better understand the phylogeographic relationships between the different strains. The neighbour-joining tree representing this phylogeographic reconstruction falls into six well-defined clusters, varying in size, both with respect to number of individuals and polymorphic positions (Figure 1; Table 1; Figure S1; Table S3).

**Figure 1.**
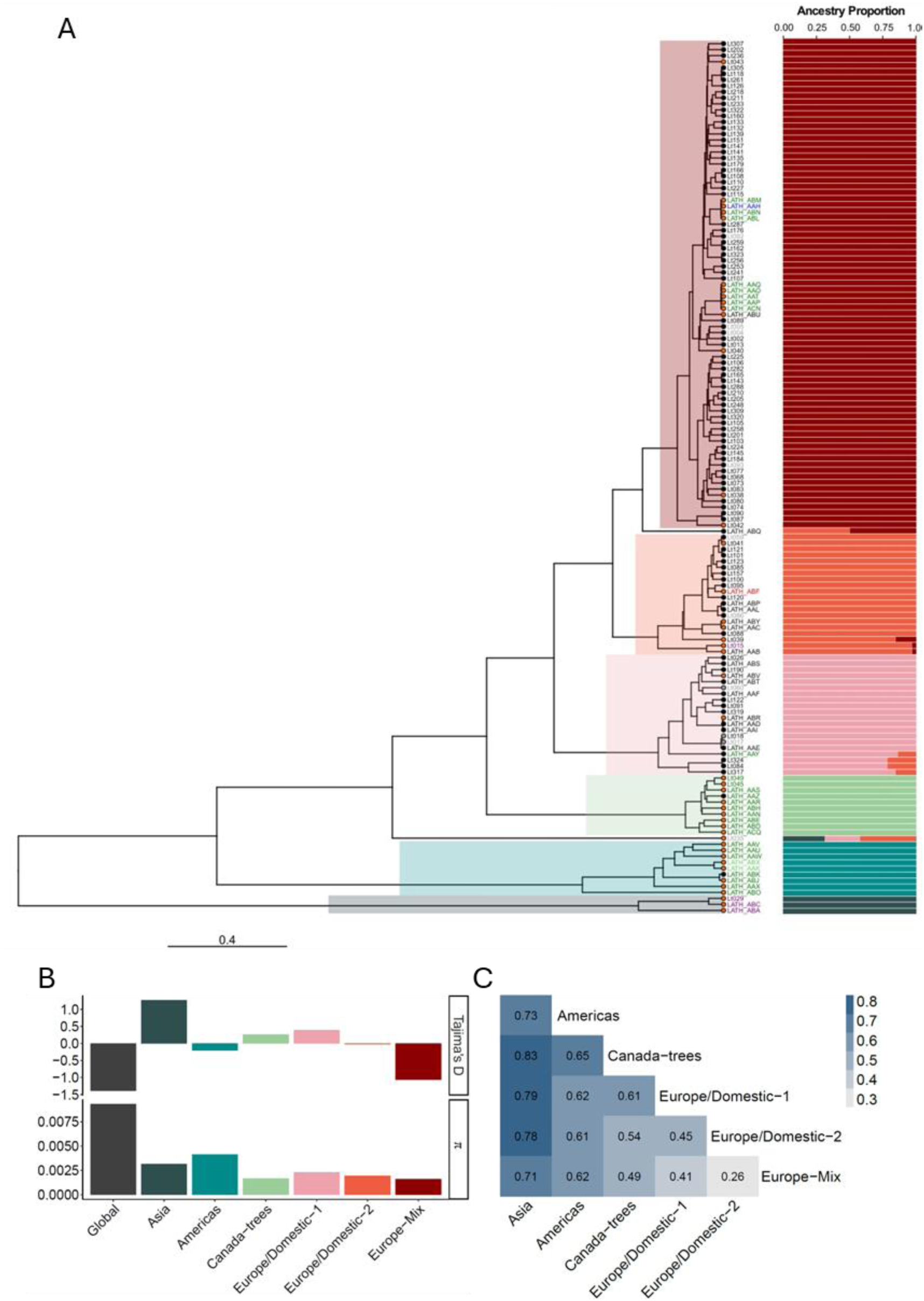
Population structure and diversity indices. **A)** Phylogenetic relationships between the 145 strains of *L. thermotolerans* and population structure (k=6) based on the SNPs dataset identified. Tip-points colour: black, anthropized; orange, wild strains. Strain name colour: black, Europe; dark green, North America; light green, South America; blue, Australia; red, Africa; purple, Asia. In light grey, no data regarding isolation source or location are disponible. Population structure colours: Asia cluster: dark green; Americas: blue; Canadian-trees: light green; Europe/Domestic 1: pink; Europe/Domestic-2: orange; Europe-mix: brown. **B)** Tajima’s D and Pairwise Nucleotide Diversity (colour codes are the same from panel A, whole-dataset values are shown in dark grey). **C)** Weir and Cockerham estimator (Fst) values across different clusters.

Population structure and admixture patterns constructed with the whole SNP dataset reveal a high degree of differentiation between strains from different environments. Six well-delimited genetic clusters (Figure 1, Figure S1-S2) are supported by both fastStructure and Admixture models, all consistent with the groups delineated by the phylogeographic relationship (Figure 1). Analysis of individual admixture patterns retrieved from the six-cluster structure of the phylogenetic analysis suggested very few admixture events in the studied dataset (Figure S2).

Using the model that integrates six clusters, the most ancestral population is composed of wild strains isolated in Asia. Subsequently, American strains segregated into two distinct groups: Americas and Canada-trees. Then, European strains appeared, and although they are isolated from similar locations and environments, the admixture patterns within this group are well resolved, revealing significant differences in population structure. Therefore, three distinct clusters can be delineated within the European group: Europe/Domestic-1, Europe/Domestic-2, and Europe-Mix, revealing not only a geographical difference between the non-European and European clusters but also an ecological difference. The non-European clusters include wild strains from non-traditional wine-producing areas, while the European clusters predominantly consist of strains isolated from anthropized environments in the Mediterranean basin, the most traditional wine-producing region. Additionally, alongside strains isolated in traditional wine regions, there are isolates from emerging wine-producing regions, such as South Africa and Australia.

The North America strains, although all isolated in wild environments, are classified into three distinct genetic clusters: Americas, Canada-trees, and Europe-mix. While the Americas cluster includes strains from both North and South America, the Canada-trees cluster predominantly encompasses strains from Canadian forests. A few strains from the Europe-mix cluster have also been isolated from Canadian forests have a genetic heritage similar to similar to European wine-related strains.

The highly structured population observed in *L. thermotolerans* is a consequence of nucleotide diversity derived from the evolutionary trajectory of the species. The mean pairwise difference (π) and Tajima’s D, which measures the difference between π and the proportion of segregating sites, indicate that the species has undergone an evolutionary pressure (Figure 1, Table S3). These forces led to a high genetic diversity in the different clusters. Concerning the entire population, the average pairwise difference between strains is 9.33e-3 bp^-1^ and Tajima’s D, -1.39. The diversity values for each cluster differ significantly from the global data, especially in the most divergent clusters, which exhibit significantly lower diversity values. This trend is not only conserved within different clusters, but also evident along the genomes (Figure S2). Clusters grouping wild strains show a significantly higher pairwise difference (4.15e-3 bp^-1^) and Tajima’s D (1.27) values compared to those grouping strains from anthropized environments (π = 1.62e-3 bp^-1^; Tajima’s D = -1.07). The presence of rare alleles and high pairwise differences, particularly within European clusters, impacts other genetic diversity indices, such as the Weir and Cockerham estimator (Fst). The Fst values in the described clusters highlight significant differences, with Asian strains showing the highest values compared to more recently differentiated clusters, such as Europe/Domestic-2 and Europe-Mix, which consist mainly of isolates from winemaking environments (Figure 1).

The phylogeographic relationships between the clusters highlight the influence of the winemaking environment on the population dynamics of *L. thermotolerans*. To determine the phylogeny within the species, we included all the *Lachancea* species, with *K. lactis* serving as an outgroup. A total of 153 orthologs for which a representative was found in each considered species was used for phylogenetic reconstruction (Table S4). The resulting phylogenetic tree, including the entire *Lachancea* clade along with 20 strains of *L. thermotolerans* representative of the species genetic diversity, corroborates the previously identified genotypic clusters (Figure 2, Figure S3). The phylogenetic relationships align with the previously described clusters, suggesting that the oldest *L. thermotolerans* strains originate from natural environments, organized according to geographical origin, while the more recent strains are predominantly associated with winemaking environments in the Mediterranean basin.

**Figure 2.**
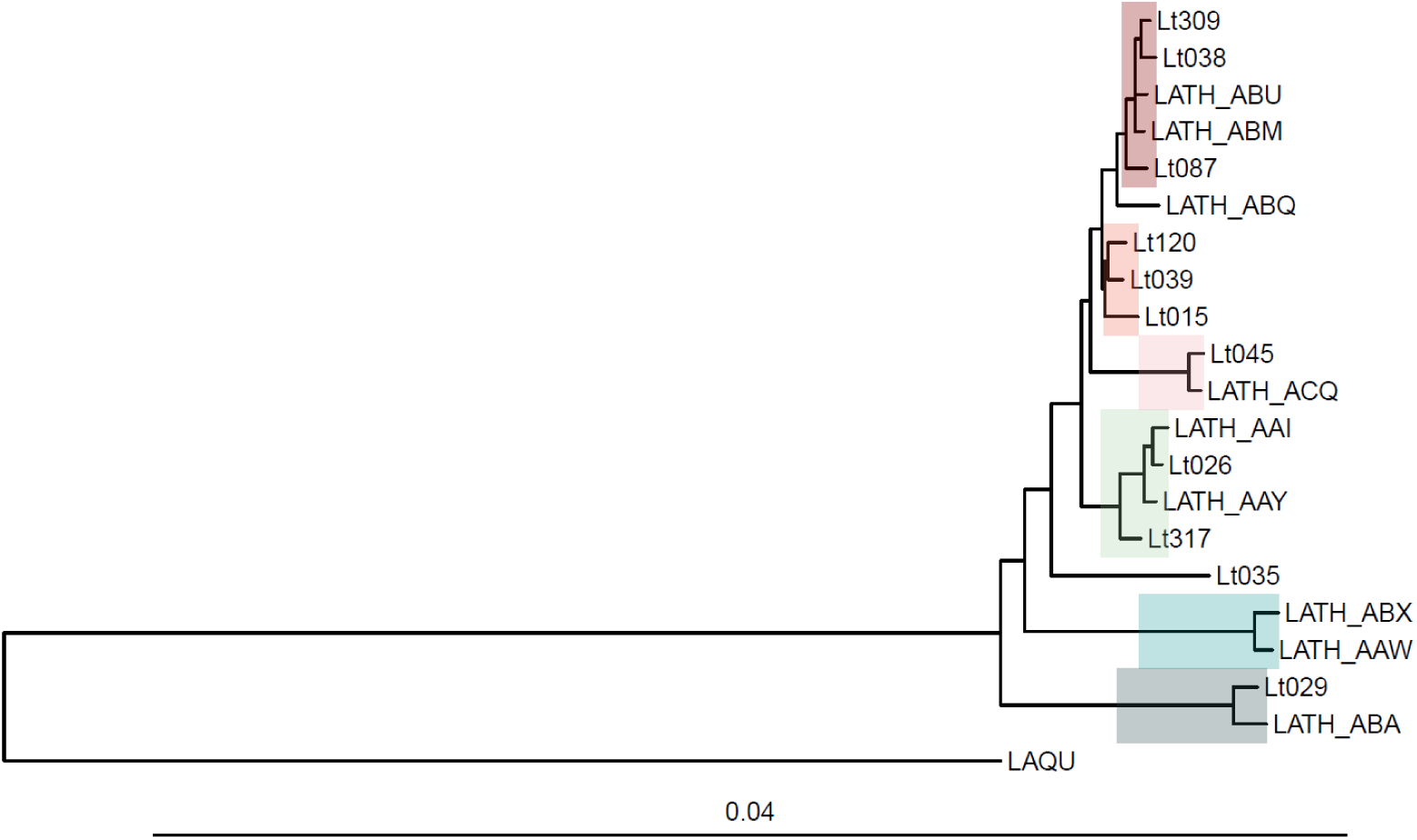
Phylogenetic reconstruction of *L. thermotolerans. L. quebecensis* has been included as outgroup. The colour code for the clusters is consistent with that used in Figure 1b.

### Wild strains of *L. thermotolerans* exhibit greater plasticity with respect to gene content

The evolutionary forces experienced by a species during its evolutionary trajectory shape not only the genetic variant content but also the gene content. Gene content is associated to the phylogeographical structure through adaptation, either through the acquisition of new genes from other taxa or through purifying selection of gene. Specific adaptations to the environment could be defined by the gain or loss of genes or by the modification of their copy number.

We used the dataset of *L. thermotolerans* strains to determine the pangenome of this species. We classified the set of genes detected in this population in four different categories based on their frequency: core and soft-core, representing high-frequency genes; shell, representing medium frequency genes; and cloud genes, representing low-frequency genes, as they are present in less than 5% of the sample.

The final gene dataset for the 145 strains of *L. thermotolerans* included 5,335 genes, with 4,948 core genes and 99 soft-core genes. Of the remaining 288 genes, 69 were classified as shell genes and 219 as cloud genes (Table S5 and S6). The species presents an open pangenome according to Heaps’ law (α = 0.56; intercept = 13.63), which may reach saturation in 5,587 genes (Figure 3A). Thus, the pangenome openness is a consequence of the content in supplemental genes that may contribute to the phenotypic variations exhibited by the strains due to the adaptive processes to specific environments. Variation in gene content helps discriminate subpopulations according to previously demonstrated population structure (Figure 1; Figure 3B). Comparing the number of different genes between each pair of clusters makes it possible to differentiate subpopulations based on their gene content. The Americas and Asia clusters have the highest number of unique genes compared to the other clusters, differing by 186 genes between them (approximately 3.5% of the total pangenome size) (Figure S4). The content of additional genes is influenced by the origin of the strains, with those coming from wild environments showing a higher number of supplemental genes (p=6.8e-3) (Table S6, Figure S4).

**Figure 3.**
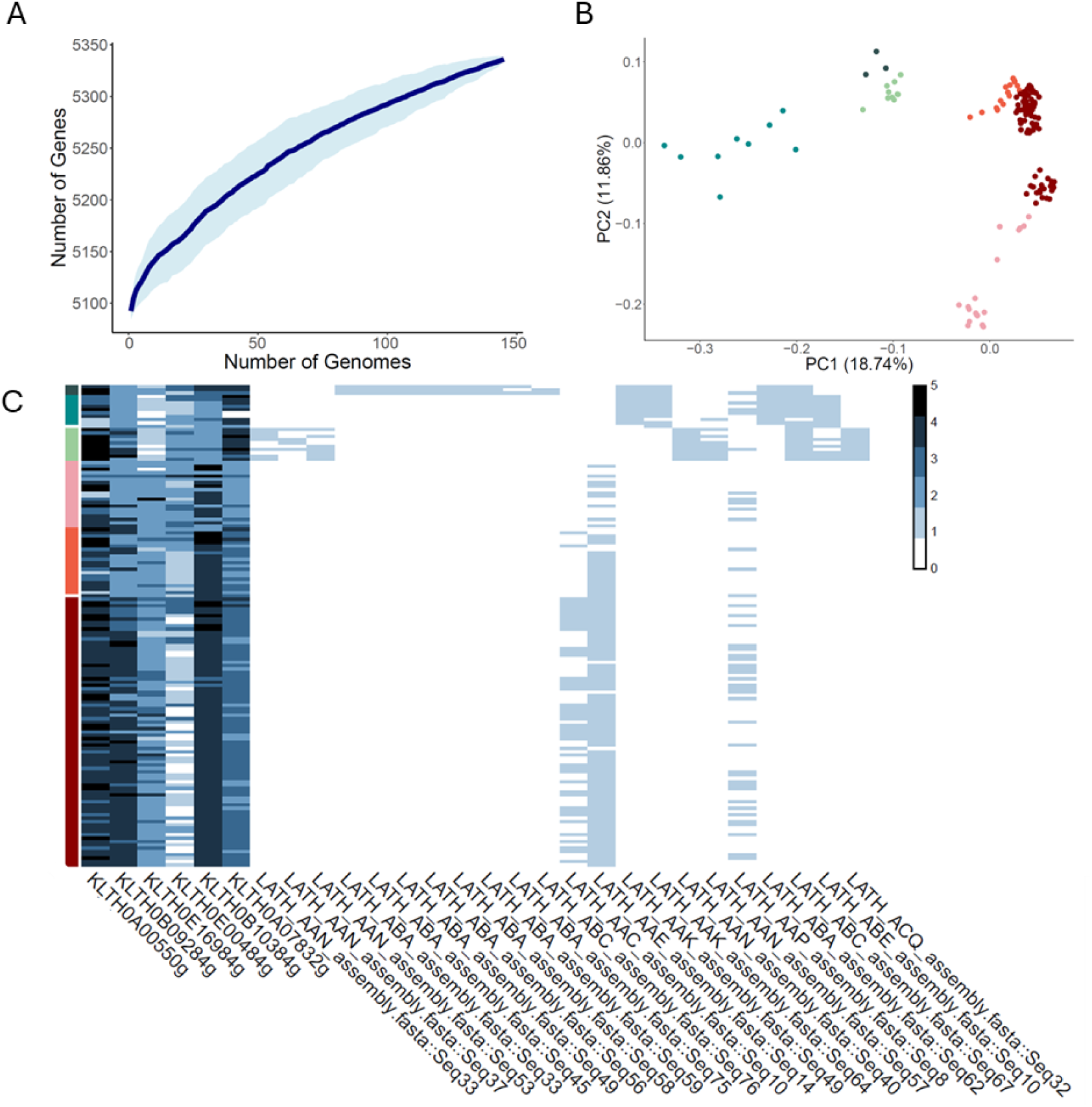
Pangenome description. **A)** Gene content as number of genomes is increased adjusted to Heaps’ law. **B)** PCA regarding gene content of each strain code. **C)** Gene content of each strain regarding high copy number genes and selected non-reference genes. The colour code for the clusters is consistent with that used in Figure 1b. In Figure 3C, blue scale indicates the number of copies per gene.

Although the acquisition or loss of genes is associated with the gain or loss of specific functions, variations in genes copy number greatly influence strains characteristics. This constitutes a common mechanism in the adaptative process to specific environments. Among the 5,092 annotated protein-coding genes in the reference, a total of 3,031 genes exhibited no change in copy number changes in all strains studied, while 2,061 genes experienced either a loss or a gain. Wild strains exhibit a significantly higher mean number of genes with copy number variations, including both decreases (p=1.4e-6) and increases (p=8.2e-3) (Figure S4). Despite this, the largest differences are observed in high copy number genes, some of them linked to specific metabolic traits (Table S7), and several showed a significantly higher number of copies in the anthropized strains than in the wild strains (Figure 3C). Among them, we highlight: KLTH0B10384g (p=8.9e-4), homolog of the *DAL5* gene in *S. cerevisiae*, which encodes an allantoin permease sensitive to nitrogen catabolite repression; KLTH0B09284g (p=2.8e-2), an ortholog of *PUG1*/*RTA1* genes in *S. cerevisiae* and member of the fungal lipid-translocating exporter (LTE) family of proteins; and KLTH0E16984g (p=5.0e-5), a high-affinity maltose permease homologous to *MAL31* in *S. cerevisiae,* whose copy number is significantly increased in Europe-mix cluster.

Among the non-referenced genes, significant differences are also observed. The content of accessory genes is one of the main drivers for strain differentiation (Figure 3B). The Americas cluster has the largest number of gained genes (119), most of which are specific of it (Figure S4, Table S6). The Americas, Asia, and Canadian-trees clusters, which appear to be the oldest within the *L. thermotolerans* species, exhibit markedly different non-reference genes content, which is cluster specific. Conversely, European clusters, associated with winemaking environments, display a similar profile, with greater similarity observed as genetic distance decreases (Figure 3B).

The supplemental genes have a different distribution in different strains and clusters. Two categories of supplemental genes among the entire dataset can be distinguished: medium frequency (as part of shell genome) and low frequency (as part of cloud genome). Non-reference genes of medium-frequency (9.85%) are much less common than those of low-frequency (90.12%). The Europe/Domestic-2 and Europe-mix clusters present several non-reference genes related to different nitrogen assimilation pathways, such as homologs of *CHA1* (Catabolic L-serine/L-threonine deaminase) and *ATO3* (Ammonia Transport Outward). Non-European clusters (i.e., Americas, Asia, and Canada-trees) have several common genes related to nitrogen metabolism. Nevertheless, some genes are cluster specific, such as those linked to aldehydes detoxification (from lignocellulose degradation) and to alternative nitrogen pathways in Canadian cluster, or those associated with iron and galactose metabolism in strains from the Asian cluster (Figure 3C, Table S8).

### Phenotypic diversity in *L. thermotolerans* is linked to the evolutionary history and the origin of subpopulations

The anthropization of the environment has impacted the evolutionary process of *L. thermotolerans*, as evidenced by the phylogenetic tree. Thus, we characterized the entire collection of strains by focusing on traits of interest from both an ecological and industrial point of view. Additionally, the interest of this species regarding its application in the wine industry is not only in its fermentative capacity, but also for the production and metabolism of organic acids, allowing the regulation of pH throughout the fermentation process. Thus, our study includes, not only the phenotypic performances of the different strains under winemaking conditions, but also other parameters of ecological interest, regarding different carbon and nitrogen sources as well as temperatures. This phenotypic characterization as well as the evaluation of the fermentation performances of the strains, determining their impact on the chemical profile of the resulting wines in laboratory-scale fermentations, can provide information on the adaptation from which the species has suffered along its evolution, niche specialisation and anthropization.

Regarding the phenotypic characterization, lag phase, maximum growth rate and proliferative efficiency (as the difference between final and initial cellular density) were extracted from each growth curve. All strains showed a good proliferation efficiency under different carbon and nitrogen sources showing no differences between different clusters and origin (Figure S5-6, Table S9). Some differences are observed in specific media (Figure 4), such as 7% ethanol, where wild strains show an increased lag phase (Wilcoxon rank-sumtest, p=3.5e-2) and a lower growth rate (p=1.7e-3) compared to the anthropized strains. These differences are also observed in the different clusters, showing the most recent ones a higher fitness (lower lag phase and higher maximum growth rate) under this condition. Some differences are also observed in other conditions, especially those regarding temperature performance. Wild strains show a lower lag phase at lower temperatures (15°C) and a lower growth rate at higher temperatures (37°C), leading to superior performance at low temperatures. Wild strains demonstrate reduced fitness, attributed to a lower maximum growth rate when using alternative carbon sources such as glycerol (p=6.7e-10) and maltose (p=1.9e-2). Across the full range of oenological conditions tested, all strains exhibited similar responses, demonstrating optimal performance under various wine fermentation conditions. Nevertheless, significant differences appear not only at high temperatures, where wild strains show a decreased efficiency (lag phase p=1.9e-3), but especially with regard to resistance to the most usual winemaking antimicrobial, sulphur dioxide. Anthropized strains show better fitness under the presence of sulphur dioxide, showing a significantly higher maximum growth rate (p=4.9e-6) and a reduced lag phase (p=2.9e-7). As the origin of the clusters becomes more recent, there is a noticeable increase in resistance to this antimicrobial, evidenced by reduced lag phases and higher growth rates. Copper is another antimicrobial commonly present in grape must as it is used in viticulture as control agent. Wild strains show a significantly higher growth rate (p=7.4e-5) compared to anthropized ones. Nevertheless, this trait is linked to specific clusters, such as European ones, showing greater fitness when this antimicrobial is present. The Canadian isolates are present in two different clusters (Canadian and Europe-Mix), which are genetically distant. Two phenotyping parameters serve as definitive factors to distinguish the two groups: copper resistance, which is exclusively associated with wild strains, and sulphur dioxide resistance, which is linked to strains genetically similar to the European-mix cluster.

**Figure 4.**
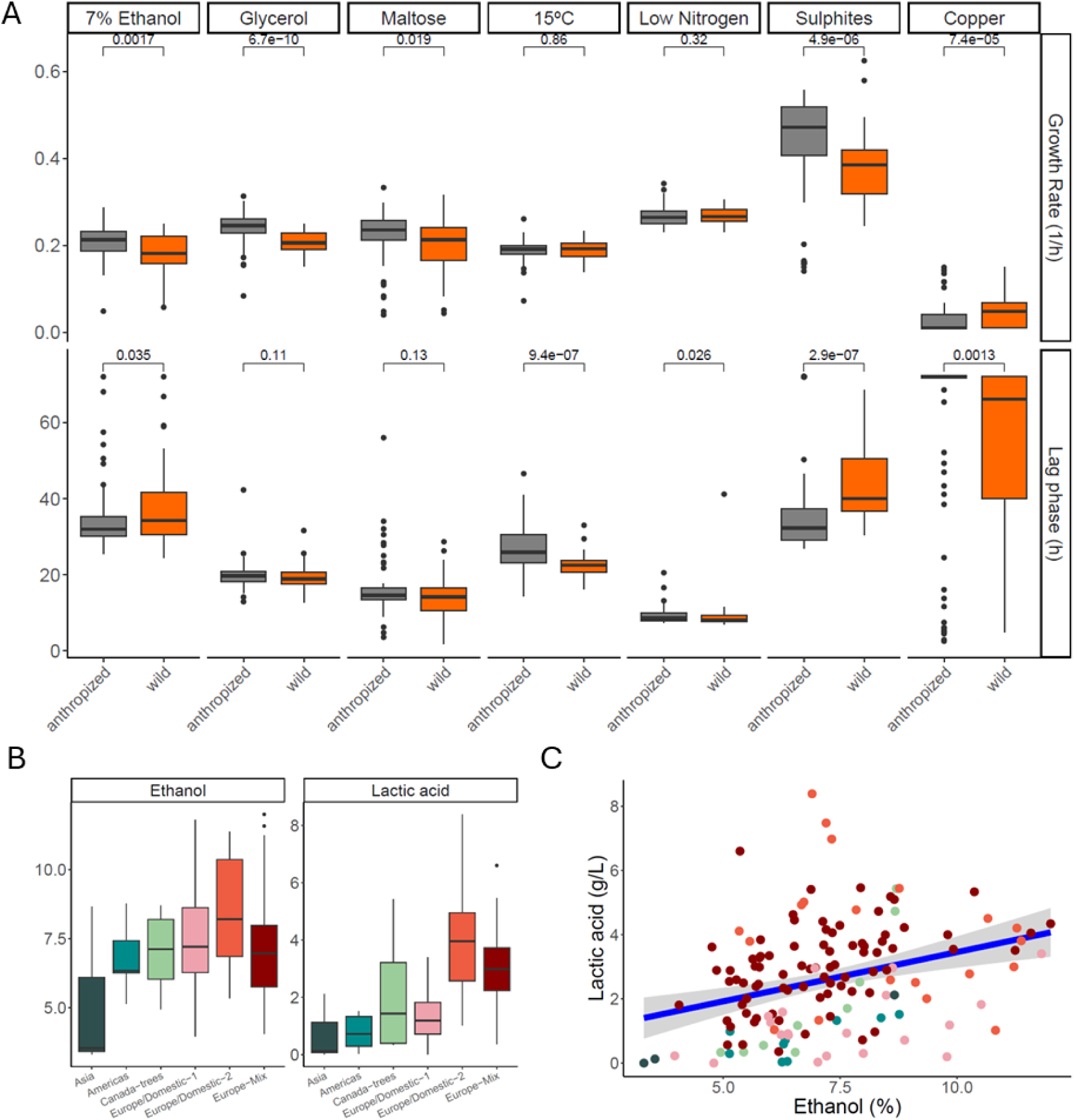
Phenotypic characterization of the strains. **A)** Phenotypic performance of the wild (orange) and anthropized (grey) groups (43 and 99 strains respectively) in the conditions showing statistical significance (Wilcoxon rank test). **B)** Ethanol and lactic acid production by the defined clusters under fermentative conditions. **C)** Ethanol and lactic acid production correlation. The colour code for the clusters is consistent with that used in Figure 1b.

The fermentative profile of the strains studied during laboratory-scale fermentations revealed important traits linked to anthropized strains (Figure 4, Figure S7, Table S10). Among the different parameters analysed, two stand out. The anthropized strains are characterized by their fructophilic character, leaving significantly lower fructose levels (p = 2.4e-2), and impacting the quantity of residual sugars (p = 4.7e-2) −glucose is similarly consumed by both groups (p=9.4e-2)−. The level of residual sugars varied from 58.54 to 51.23 g/L for wild and anthropized strains, being significantly high for the Asian strains (98.96 g/L) and low for the European-Mix strains (48.16 g/L). Another important trait of the anthropized subset of strains is its high lactic acid production, which shows a significant correlation with ethanol production (Rho=0.369; p-value < 2.2e-16) (Figure 4). Lactic acid production by the anthropized strains reaches a significantly higher mean production of 2.98 g/L compared to the wild subset of strains (1.88 g/L; p = 1.7e-4). Important differences are observed regarding specific clusters, varying its mean production from 0.75 to 3.98 g/L for the Asian and Europe/Domestic-2 clusters.

To determine the relationship between the phylogenetic relationships of the strains and the presented phenotypic landscape of each strain, we determined the correlation between the two datasets. We used the constructed neighbour-joining tree to establish the cophenetic distance between each pair of strains, thereby determining the phylogenetic distance between them. At the same time, the phenotypic differences of each pair were assessed based on the Euclidean distance regarding both the phenotypic characterization and the oenological performance of strain (Figure S8, Table S11). We found a significant positive correlation between the two distances (Rho=0.441; p-value < 2.2e-16) (Figure S8). Moreover, 57 out of the 63 phenotypic traits analyzed in this study showed a significant phylogenetic signal (Pagel’s λ p < 0.05; Figure S8, Table S12), suggesting a connection between phenotype and phylogeny.

## DISCUSSION

Anthropization is a major selective force arising from the environmental conditions imposed by human activities. This adaptation involves different forms of genomic reorganization that enable microorganisms to rapidly adapt to imposed environmental challenges (Villarreal et al., 2024). Despite *S. cerevisiae* being the most studied eukaryotic species due to its industrial relevance and its establishment as a model species for evolutionary and niche adaptation studies in yeasts, expanding attention to studying different genera and species could allow a better comprehension of global patterns of adaptation to the anthropic (and fermentative) environment experienced by yeasts. Analysing the genotypic and phenotypic diversity and the main adaptative traits in fermentative yeasts will provide a better understanding of the evolutionary dynamics of nature and the association with anthropic environments (Villarreal et al., 2022).

In our work, we focused on *L. thermotolerans,* a non-*Saccharomyces* yeasts species, involved in winemaking, which has gained attention in recent years due to its potential as a fermentative alternative to cope with climatic conditions (Vicente et al., 2023). Through whole-genome sequencing of a large set of 145 strains, we determined the population structure of the species, deciphering the evolutionary trajectory and confirming the influence of winemaking practices on yeasts evolution and subpopulation differentiation (Freel et al., 2014; Hranilovic et al., 2017). We identified differential gene content related to this adaptation process that is linked to different phenotypes. Extensive phenotyping, including analysis of fermentation performance, revealed different behaviours among the subpopulations studied. Furthermore, the adaptation process, influencing gene content in some cases, has shaped the phenotypic landscape of the species. Additionally, our fermentation analyses demonstrate specific traits such as lactic acid production and fructose consumption in wine-related yeast strains. The study provides strong evidence for the effect of anthropization on yeast evolutionary processes and how human practices, acting as an unintended force, have once again driven microbial evolution and adaptation.

The first evidence of *L. thermotolerans* evolution and population differentiation revealed a significant influence of the geographical and ecological origin of the strains. This study shows a strong differentiation between anthropized strains compared to wild isolates, the latter presenting differentiation based on geographical origin (Hranilovic et al., 2017). Our results, based on SNPs, confirm this distribution, revealing three different clusters of wild strains originating from different regions (Americas, Asia, and Canadian-trees), as well as three additional clusters of anthropized strains, mainly originating from Europe (Europe/Domestic-1, Europe/Domestic-2, and Europe-mix). All subpopulations exhibit a distinct differentiation pattern with few admixture events and high fixation indices. Despite the species shows high genetic diversity (π, 9.33e-3 bp^-1^), as a consequence of the different and well-differentiated clusters, an abundance of rare alleles is also observed (Tajima’s D, -1.39), suggesting a currently expanding population. The lower genetic diversity, as well as Tajima’s D values near 0, indicative of recent bottlenecks, in the anthropized clusters, suggest influence of purifying selection. Comparative analysis reveals high diversity values in related species, such as those in the *Kluyveromyces* clade undergoing domestication processes, like *K. lactis* or *Kluyveromyces marxianus,* with average pairwise diversity values of 2.8e-2 and 1.2e-2 bp^-1^, respectively. However, other species exhibit significantly lower values, such as *S. cerevisiae,* with values around 3e-3 bp^-1^ (Peter et al., 2018). The values observed in species undergoing strong selective process (i.e, *S. cerevisiae*) are similar to those observed in *L. thermotolerans* clusters from anthropized environments (around 2e-3 bp^-1^).

The genetic distance (i.e, fixation index) between different clusters, together with the similarity shown in the phylogenetic relationship with *K. lactis*, suggests that the Asian cluster is both the most isolated and the most ancestral subpopulation. These data imply that this cluster could be the first to diverge, and that, as in *S. cerevisiae*, species evolution probably initially followed geographical patterns and then niche adaptation, with the European isolates influenced by winemaking practices being the most recent. Thus, anthropization appears to be the main driver of species evolution and genetic differentiation. However, this anthropization is likely not a single event but rather a combination of different adaptation processes, as evidenced by the presence of well-differentiated clusters.

As observed in other species, strong evidence of anthropization and adaptation to wine environment by modifying gene content is found in *L. thermotolerans*. Gene content analysis improves understanding of the evolutionary process in a species, as gradual changes that accumulate over time become more evident (Silva et al., 2023). Variation in gene copy number, along with introgression and horizontal gene transfer, have been described as among the most common genetic adaptations, serving as the primary mechanisms of adaptation to the wine environment (Marsit & Dequin, 2015; Steenwyk & Rokas, 2018). Purifying selection is observed in anthropized strains, with fewer genes showing copy number alterations as well as supplemental genes. However, important genes typically associated with anthropization in *S. cerevisiae*, such as those related to the assimilation of alternative carbon sources (Peris et al., 2023) or nitrogen metabolism (García-Ríos & Guillamón, 2022), exhibit differences in the studied anthropized strains of *L. thermotolerans*, particularly in homologous genes to *DAL5* and *MAL31*.

Nitrogen acts as a limiting nutrient in most of wine fermentations due to its scarcity in natural grape must. In *S. cerevisiae,* oligopeptides are usually transported by carrier proteins Ptr2 and Dal5, responsible for the transport of di/tripeptides and dipeptides, respectively. Dal5 constitutes the predominant dipeptide transporter in some *S. cerevisiae* strains, despite its greater affinity for molecules other than for peptides (Becerra-Rodríguez et al., 2020). Similarly, despite the low concentration of maltose in grape must, *MAL3* locus is primarily duplicated among wine *S. cerevisiae* isolates. It is hypothesized that this duplication could potentially facilitate the assimilation of secondary substrates or provide other functions conferring an adaptive advantage (Steenwyk & Rokas, 2018). Thus, increased copy numbers of some genes (i.e., *DAL5* and *MAL31*) in *L. thermotolerans*, likely play a role in improving the fitness of these isolates in this environment, facilitating enhanced peptide uptake and the assimilation of alternative carbon sources by anthropized strains, respectively. This fact is confirmed when phenotyping data are analysed, as strains from winemaking environments show higher fitness when using maltose as sole carbon source, suggesting the importance of gene dosage and/or acquisition in *L. thermotolerans*.

Similar to other anthropized yeasts species such as *S. cerevisiae* and *L. cidri,* we find other relevant phenotypic traits regarding niche adaptation in anthropized *L. thermotolerans* strains, such as increased respiration (higher fitness assimilating glycerol) and fructose consumption, as well as increased ethanol and sulphite tolerance (Onetto et al., 2023; Villarreal et al., 2024; Warringer et al., 2011).These genomic and phenotypic features suggest convergent adaptative mechanisms to human-related environments, as phylogenetically distant yeast species linked to different fermentative processes show similar adaptations (Villarreal et al., 2024). Additionally, most of the phenotypic traits studied have a direct link to the phylogeny of the species, reinforcing the role of adaptative evolution consequence of the anthropization process, and highlighting how key functional traits of ecological and industrial relevance are linked to phylogeny and genomic traits of yeasts (Opulente et al., 2024; Ruiz et al., 2023).

Specific phenotypic traits are associated to wild strains, among which tolerance to copper and lower temperatures tolerance stand out as the most relevant. Unlike observations in *S. cerevisiae,* where the use of copper as a fungicide in vineyards led to increased tolerance in anthropic isolates (De Guidi et al., 2024; García-Ríos & Guillamón, 2019, 2022; Onetto et al., 2023), we found that wild isolates of *L. thermotolerans* from Canadian-trees cluster exhibit increased fitness in the presence of copper compared to anthropic isolates. Additionally, Canadian isolates confirm the possibility of yeast migration between different environments. We hypothesize the presence of feral strains among the Canadian isolates. Canadian strains from Europe-Mix clusters could have been associated with wine-producing environments in the past, where they underwent an adaptation process. Subsequently, they could have migrated from this location and become isolated in a different ecosystem, while retaining the genetic and phenotypic characteristics of the anthropic environment, such as sulphites resistance, without showing genotypic or phenotypic traits linked to it (e.g., copper resistance). Regarding temperature, this environmental factor, along with the substrate, is one of the main conditions defining ecological diversity and niche distribution by yeast taxa (Spurley et al., 2022). In the *Saccharomyces* genus, this trait has been defined as an ancestral trait influenced by the ecological and geographical distribution of the different *Saccharomyces* populations (Peris et al., 2023). We highlight the better performance of wild strains at low temperatures, which may improve survival in wild environments such as those associated with insects or subboreal locations (Hranilovic et al., 2017). It has been verified that colonization of bee digestive tracts by *L. thermotolerans* is associated with lower temperatures (Kogan et al., 2023).

Based on the results obtained in this study, we propose lactic acid production as a hallmark of anthropization in *L. thermotolerans.* Some studies have described increased lactic production in specific groups of *S. cerevisiae* from distilleries, sake, and rum as an adaptation to fermentation under stressful conditions, linked to an increased glycolytic flux. Although *S. cerevisiae* does not have fully active lactate dehydrogenase enzymes (unlike *L. thermotolerans*), lactic acid production is coupled to the α-ketoglutarate production from 2-hydroxyglutarate and to methylglyoxal degradation, produced from glycolytic intermediates (Monnin et al., 2024). In *L. thermotolerans,* it is generally accepted that lactic acid production may play a role in regulating the redox potential (Shekhawat et al., 2020; Vicente et al., 2021).

A recent study examining the redox balance during fermentation in high and low lactic acid-producing strains, as well as in other yeasts species, revealed a favoured ratio of NAD+ despite the lower concentration of total NAD(H) compared to other taxa. This suggests that although highly lactic acid-producing strains of *L. thermotolerans* have a lower total amount of total NAD(H), the favoured ratio to NAD+, likely derived from lactic acid production, is sufficient to maintain an adequate proportion of redox cofactors and support high glycolytic flux during the initial stages of the fermentation (Tyibilika et al., 2024). Previous studies have shown that low- and high-producing strains are phylogenetically separated (Battjes et al., 2023; Gatto et al., 2020). Additionally, it has been demonstrated that increased transcription of *LDH2* (Gatto et al., 2020; Sgouros et al., 2020), which has specific transcriptional factors linked to low-producing strains, is associated with higher lactic acid production (Battjes et al., 2023).

In this study, we have demonstrated the link between lactic acid production and the evolutionary trajectory of the species, as this trait generally is associated with anthropized strains that are more distant from *K. lactis*. The fermentative conditions imposed by the winemaking environment can lead to an increase in glycolytic flux, modifying the redox cofactors ratios. While the Whole Genome Duplication (WGD) that occurred 100 million years ago allowed *S. cerevisiae* to have 5 different ADHs, providing flexibility to cope with increased metabolic flux (Dashko et al., 2014), *L. thermotolerans*, a pre-WGD Crabtree-positive species, shows 2 genes coding for ADHs. Therefore, it may need an alternative route to support the fermentative condition that provokes an unbalanced ratio of redox cofactors. We hypothesize that anthropized strains of *L. thermotolerans,* to cope with this condition, may have adapted their metabolism by developing or maintaining an alternative route (lactic acid production) for redox cofactor regeneration, allowing greater fitness during the fermentation process. However, further research is needed to confirm this hypothesis, particularly regarding the redox cofactor balance during fermentation, the kinetics of sugar consumption and metabolite production, as well as the regulation of the fermentation process. This entails studying a larger number of strains of *L. thermotolerans*.

Overall, the findings of this work provide valuable insight into the anthropization process in a key non-*Saccharomyces* yeast species. Studying the genetic and phenotypic diversity within this yeast has shown that fermentation processes result in similar adaptations in phylogenetically distant species, but also how specific groups exhibit unique traits to improve their fitness in these environments. Moreover, expanding our understanding of the general evolutionary dynamics in yeasts, alongside niche-specific adaptation processes, could facilitate efficient exploitation of yeasts for industrial purposes.

## CONFLICT OF INTEREST

The authors have no conflicts of interest.

## ACKNOWLEDGEMENTS

Funding for this research was provided by the LowpHWine Companies Consortia through the CDTI project LowpHWine (IDI-20210391) and the Spanish Ministry of Science and Innovation under the VinSegCalClim project (PID2020-119008RB-I00). Javier Vicente conducted this research under a fellowship from Complutense University of Madrid (CT58/21-CT59/21). Joseph Schacherer is supported by a European Research Council (ERC) Consolidator grant (ERC-CoG 772505). Joseph Schacherer is a member of the Institut Universitaire de France.

## AUTHOR CONTRIBUTION

Conceptualization: J.V., D.M., A.S. Investigation: J.V., A.F., K.F. Data analysis and visualization: J.V., A.F., J.S., S.B., A.S. Funding: J.S., S.B., D.M., A.S. Writing: J.V., A.S. All authors read and approved the final manuscript.

## DATA AVAILABILITY STATEMENT

The data presented in this study can be accessed on NCBI Bioproject: PRJNA1111406 and PRJEB29656. Accessions are given in Table S1.

## Notes

### Competing Interest Statement

The authors have declared no competing interest.

