## Supplementary figures for "Whole-genome sequencing and phenotyping reveal specific adaptations of *Lachancea thermotolerans* to the winemaking environment"

Figure S1. Population structure using 2 to 8 populations determined using FastStructure.

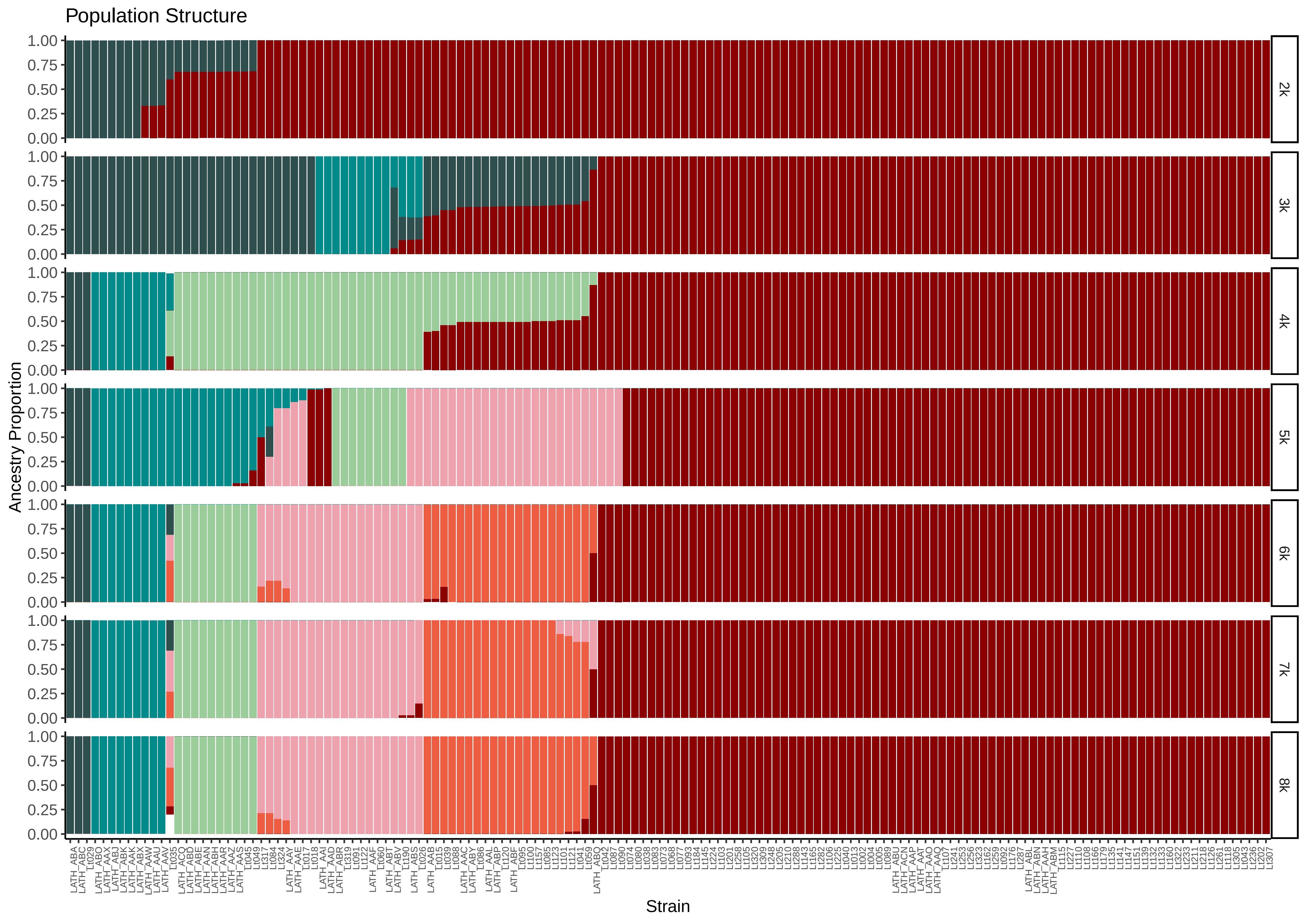
Population Structure
Ancestry Proportion
1.00
0.75
0.50
0.25
0.00
2k
3k
4k
5k
6k
7k
8k
Strain

Figure S2. A) Admixture proportions using k=6 (colour code for clusters are the same as in Figure 1a). Diversity indices along genome along the different defined clusters (colours show different chromosomes): b) Pairwise Nucleotidic Diversity. C) Tajima's D.

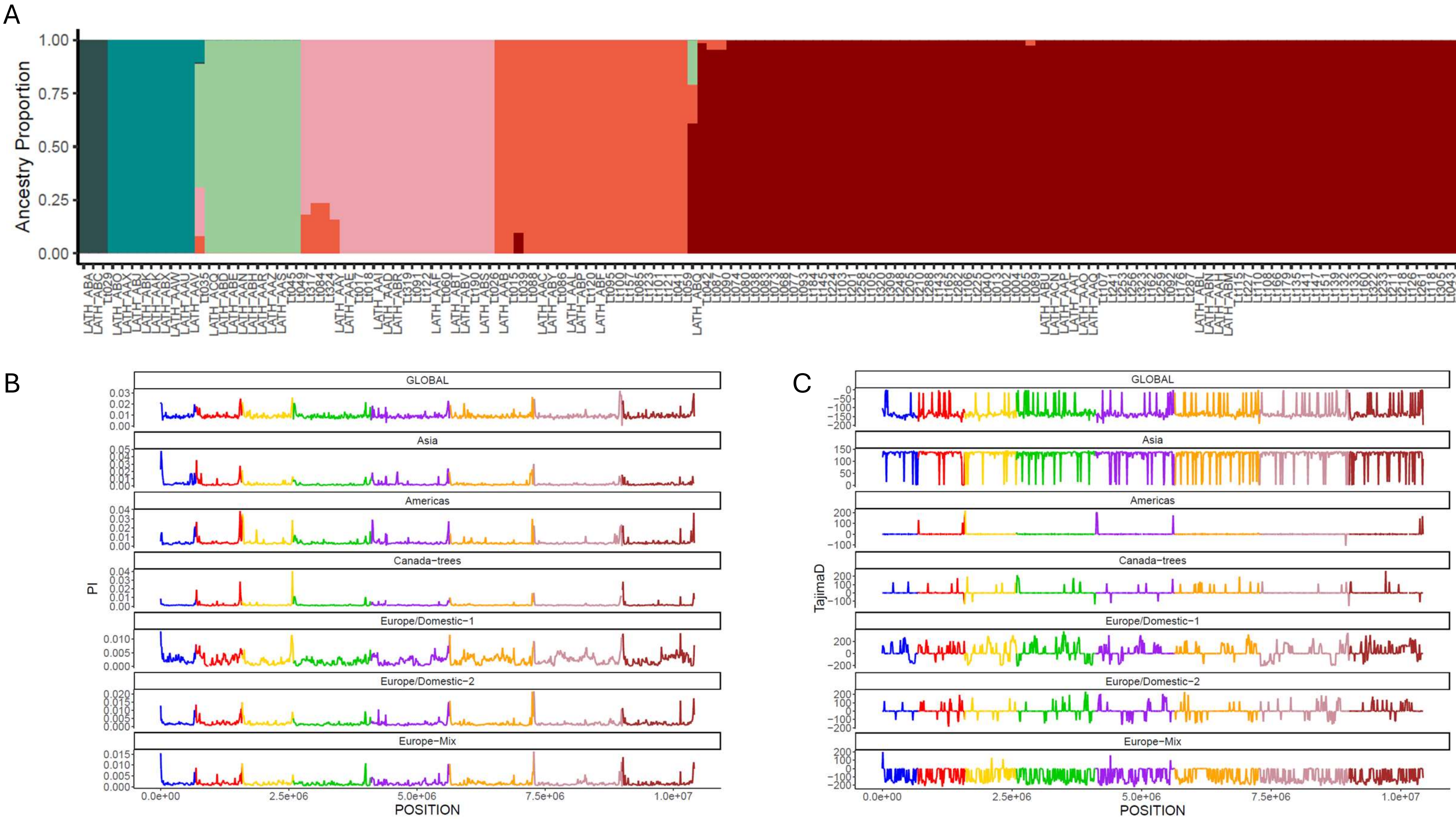
A
Ancestry Proportion
1.00
0.75
0.50
0.25
0.00
B
PI
GLOBAL
Asia
Americas
Canada-trees
Europe/Domestic-1
Europe/Domestic-2
Europe-Mix
0.0e+00
2.5e+06
5.0e+06
7.5e+06
1.0e+07
POSITION
C
TajimaD
GLOBAL
Asia
Americas
Canada-trees
Europe/Domestic-1
Europe/Domestic-2
Europe-Mix
0.0e+00
2.5e+06
5.0e+06
7.5e+06
1.0e+07
POSITION

Figure S3. Phylogenetic reconstruction of the *Lachancea* genus using *K. lactis* as outgroup (colour code for *L. thermotolerans* clusters are the same as in Figure 1a).

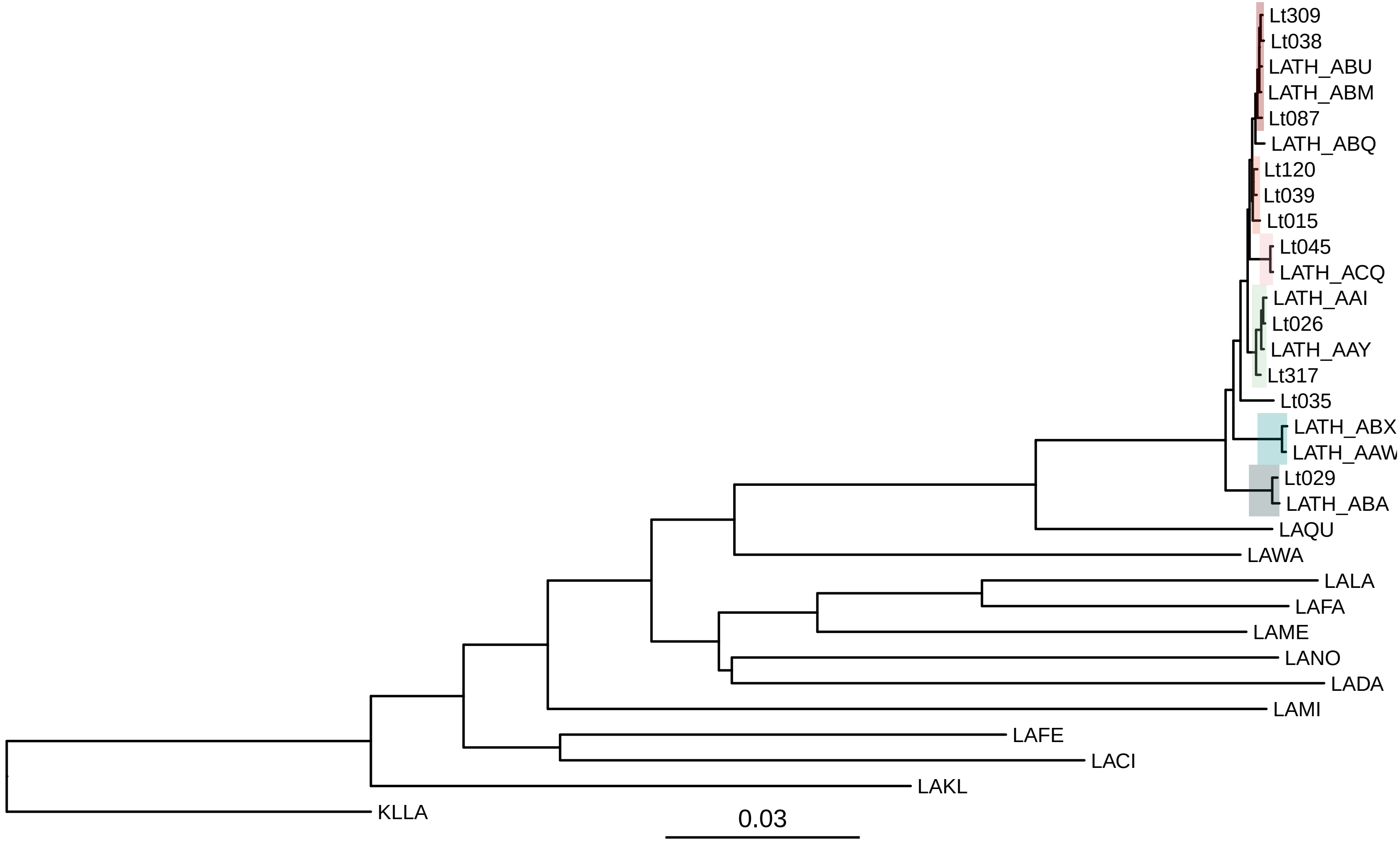
Lt309
Lt038
LATH_ABU
LATH_ABM
Lt087
LATH_ABQ
Lt120
Lt039
Lt015
Lt045
LATH_ACQ
LATH_AAI
Lt026
LATH_AAY
Lt317
Lt035
LATH_ABX
LATH_AAW
Lt029
LATH_ABA
LAQU
LAWA
LALA
LAFA
LAME
LANO
LADA
LAMI
LAFE
LACI
LAKL
KLLA
0.03

Figure S4. Gene content shared between strains (colour code for clusters are the same as in Figure 1a). A) Number of different genes between clusters. B) Number of genes showing a gain/loss in CNV in the anthropized and wild strains. C) Number of supplemental genes in anthropized and wild strains. D) Number of different non-reference genes between clusters (Wilcoxon rank-sumtest p-value is shown).

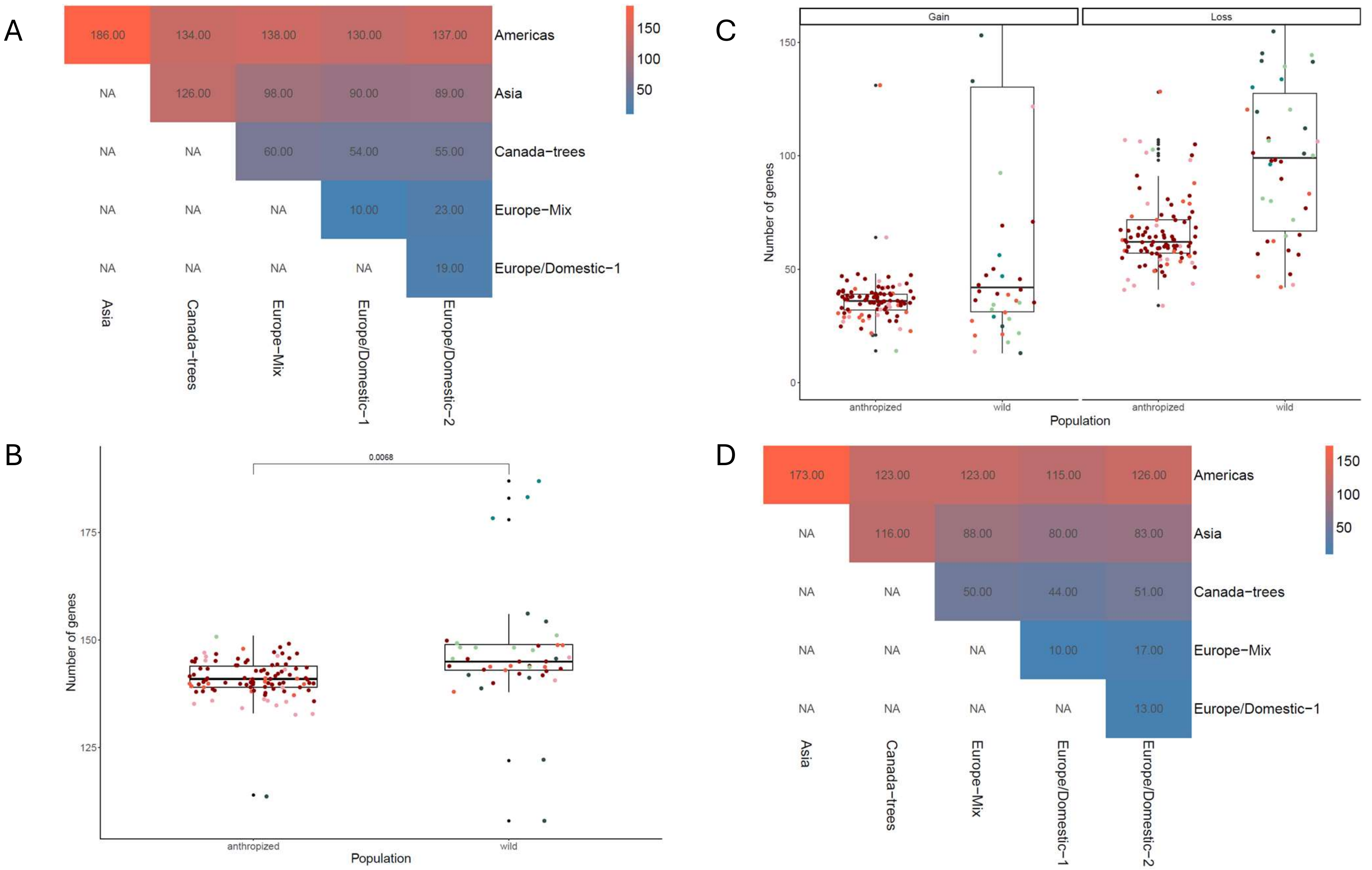
A
186.00
134.00
138.00
130.00
137.00
Americas
NA
126.00
98.00
90.00
89.00
Asia
NA
NA
60.00
54.00
55.00
Canada−trees
NA
NA
NA
10.00
23.00
Europe−Mix
NA
NA
NA
NA
19.00
Europe/Domestic−1
Asia
Canada−trees
Europe−Mix
Europe/Domestic−1
Europe/Domestic−2
150
100
50
B
0.0068
Number of genes
175
150
125
anthropized
wild
Population
C
Gain
Loss
Number of genes
150
100
50
0
anthropized
wild
anthropized
wild
Population
D
173.00
123.00
123.00
115.00
126.00
Americas
NA
116.00
88.00
80.00
83.00
Asia
NA
NA
50.00
44.00
51.00
Canada−trees
NA
NA
NA
10.00
17.00
Europe−Mix
NA
NA
NA
NA
13.00
Europe/Domestic−1
Asia
Canada−trees
Europe−Mix
Europe/Domestic−1
Europe/Domestic−2
150
100
50

Figure S5. Phenotypic characterization regarding different environmental factors. Growth rate and lag phase are shown for each media (Wilcoxon rank-sumtest p-values are shown).

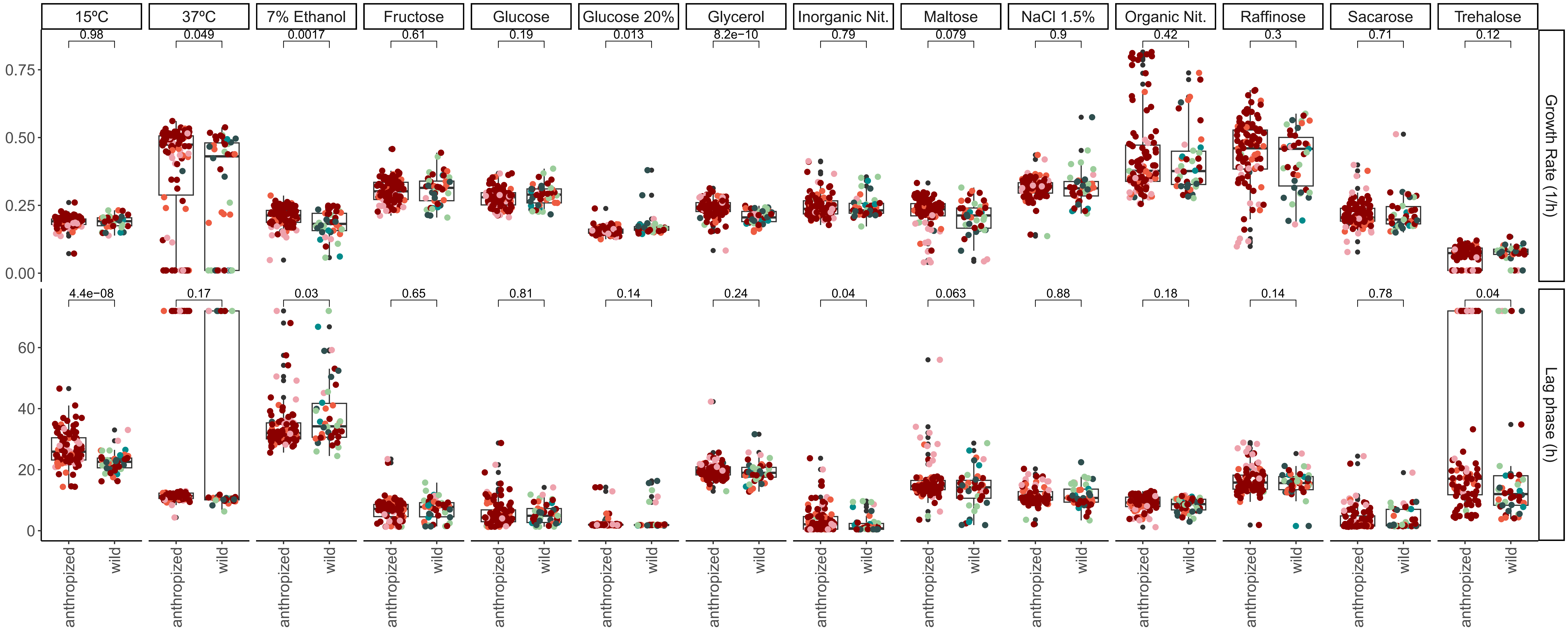

15ºC
37ºC
7% Ethanol
Fructose
Glucose
Glucose 20%
Glycerol
Inorganic Nit.
Maltose
NaCl 1.5%
Organic Nit.
Raffinose
Sacarose
Trehalose
0.98
0.049
0.0017
0.61
0.19
0.013
8.2e−10
0.79
0.079
0.9
0.42
0.3
0.71
0.12
4.4e−08
0.17
0.03
0.65
0.81
0.14
0.24
0.04
0.063
0.88
0.18
0.14
0.78
0.04
Growth Rate (1/h)
Lag phase (h)
0.00
0.25
0.50
0.75
0
20
40
60
anthropized
wild

Figure S6. Phenotypic characterization regarding different oenological factors. Growth rate and lag phase are shown for each media (Wilcoxon rank-sumtest p-values are shown).

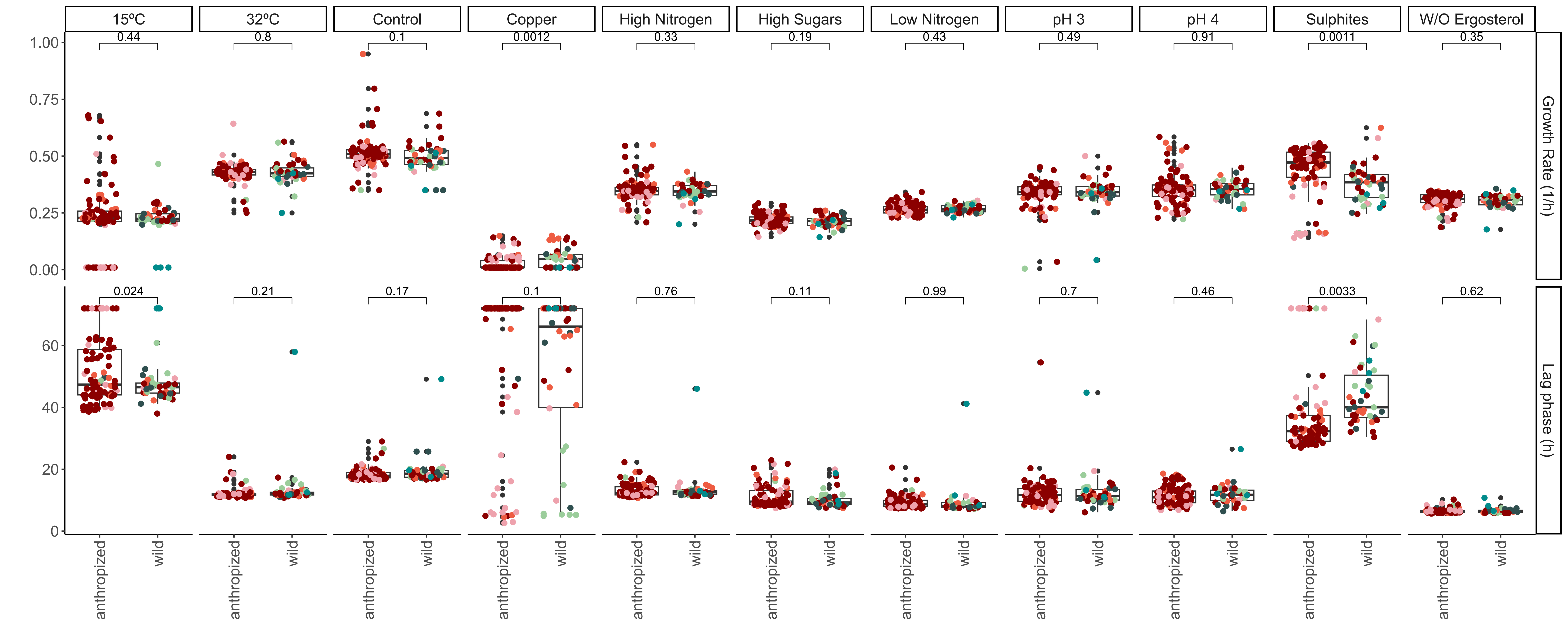

15ºC
32ºC
Control
Copper
High Nitrogen
High Sugars
Low Nitrogen
pH 3
pH 4
Sulphites
W/O Ergosterol
0.44
0.8
0.1
0.0012
0.33
0.19
0.43
0.49
0.91
0.0011
0.35
0.024
0.21
0.17
0.1
0.76
0.11
0.99
0.7
0.46
0.0033
0.62
1.00
0.75
0.50
0.25
0.00
60
40
20
0
Growth Rate (1/h)
Lag phase (h)
anthropized
wild

Figure S7. Oenological parameters at the end of the laboratory-scale fermentations in SGM (Wilcoxon rank-sumtest p-values are shown).

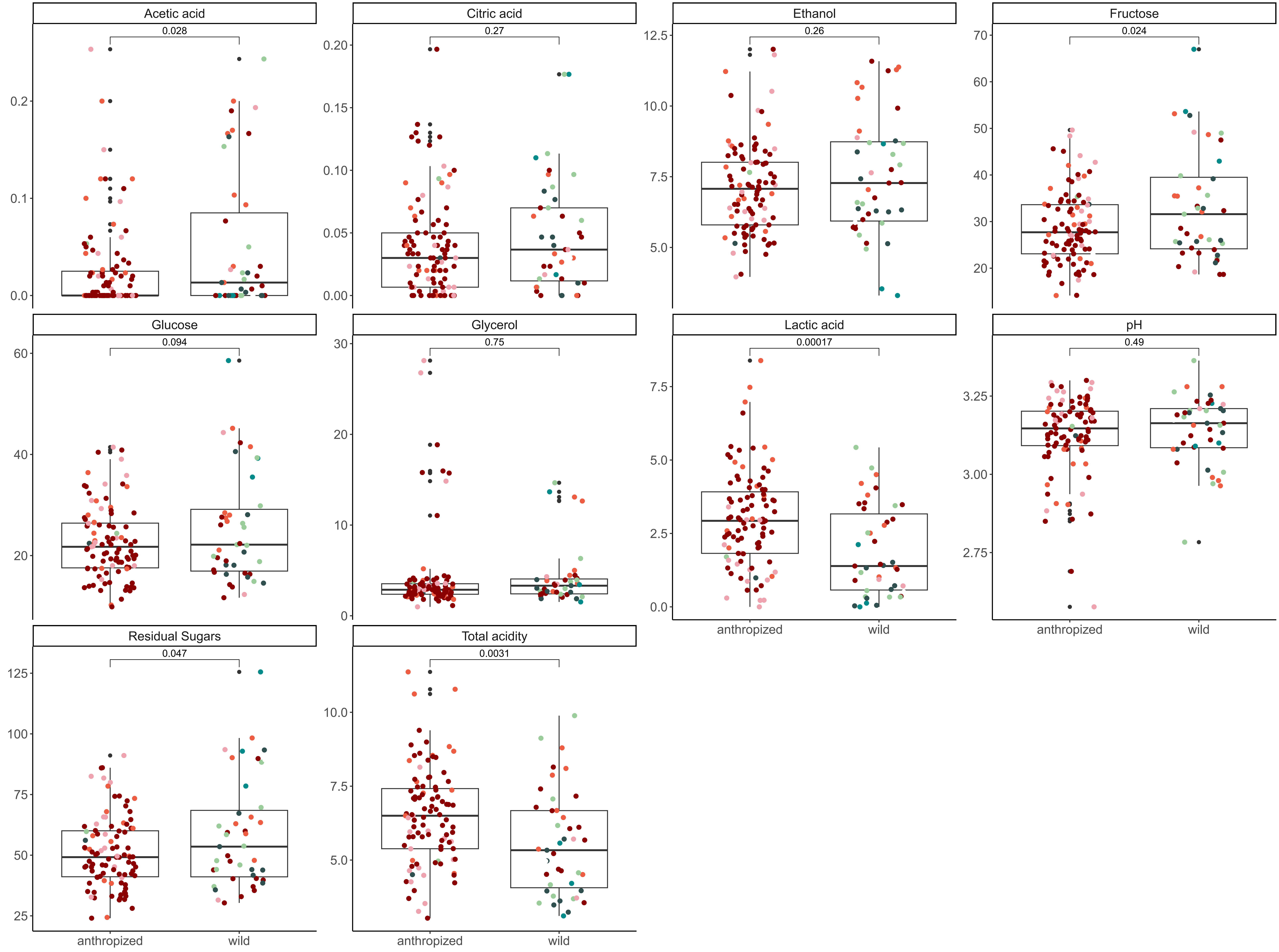

Acetic acid
0.028
Citric acid
0.27
Ethanol
0.26
Fructose
0.024
Glucose
0.094
Glycerol
0.75
Lactic acid
0.00017
pH
0.49
Residual Sugars
0.047
Total acidity
0.0031
anthropized
wild

Figure S8. Phenotype and phylogeny correlation. A) Pearson's correlation between phenotypic and phylogenetic distance (Rho=0.441; p-value < 2.2e-16). B) Phylogenetic signal of each phenotypic trait and. In grey, $p<0.05$; in red $p>0.05$.

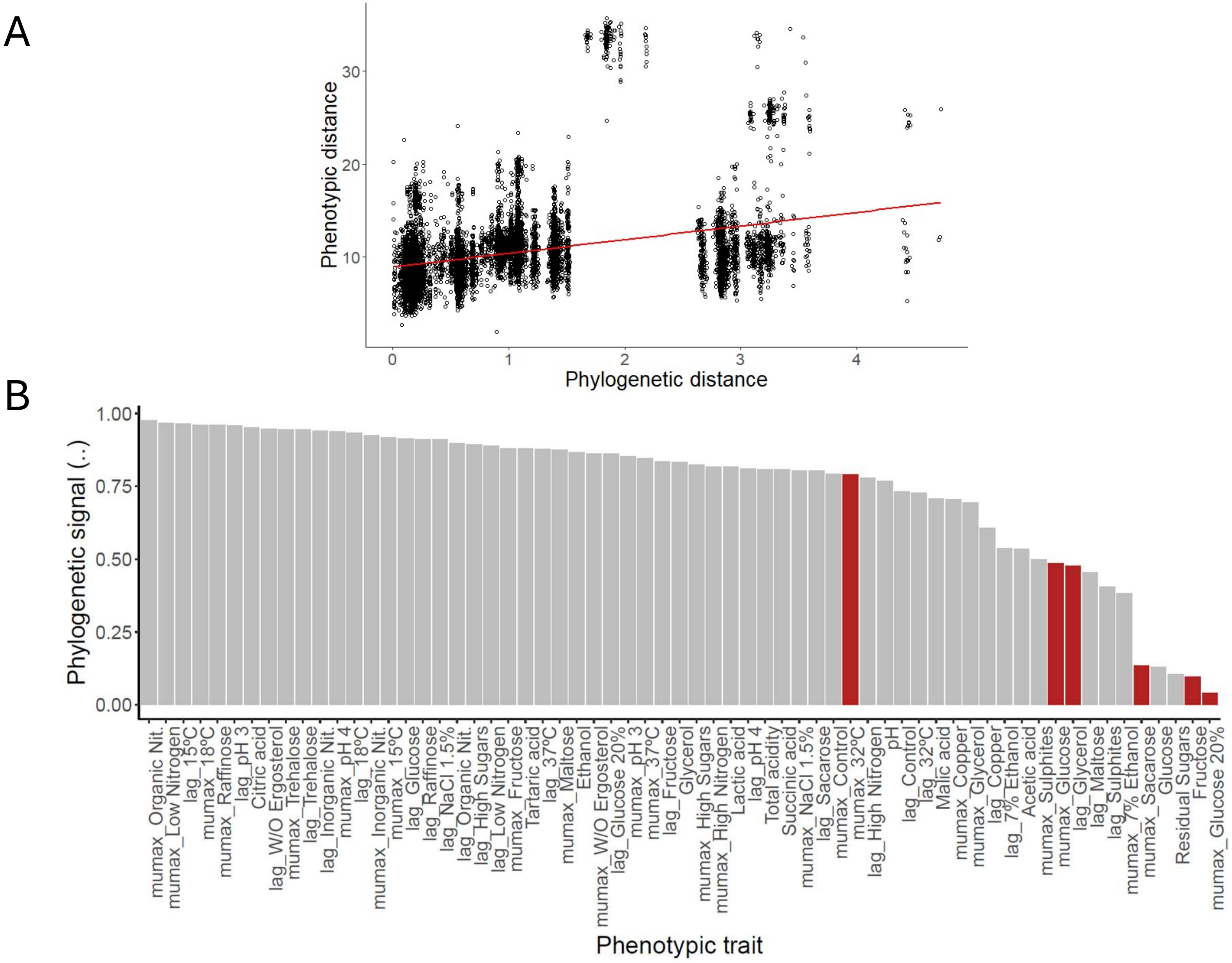

A
Phenotypic distance
Phylogenetic distance
B
Phylogenetic signal (..)
Phenotypic trait
mumax_Organic Nit.
mumax_Low Nitrogen
lag_15°C
mumax_18°C
mumax_Raffinose
lag_pH 3
Citric acid
lag_W/O Ergosterol
mumax_Trehalose
lag_Trehalose
lag_Inorganic Nit.
mumax_pH 4
lag_18°C
mumax_Inorganic Nit.
mumax_15°C
lag_Glucose
lag_Raffinose
lag_NaCl 1.5%
lag_Organic Nit.
lag_High Sugars
lag_Low Nitrogen
mumax_Fructose
Tartaric acid
lag_37°C
mumax_Maltose
Ethanol
mumax_W/O Ergosterol
lag_Glucose 20%
mumax_pH 3
mumax_37°C
lag_Fructose
Glycerol
mumax_High Sugars
mumax_High Nitrogen
Lactic acid
lag_pH 4
Total acidity
Succinic acid
mumax_NaCl 1.5%
lag_Sacarose
mumax_Control
mumax_32°C
lag_High Nitrogen
pH
lag_Control
lag_32°C
Malic acid
mumax_Copper
mumax_Glycerol
lag_Copper
lag_7% Ethanol
Acetic acid
mumax_Sulphites
mumax_Glucose
lag_Glycerol
lag_Maltose
lag_Sulphites
mumax_7% Ethanol
mumax_Sacarose
Glucose
Residual Sugars
Fructose
mumax_Glucose 20%
